# Mechanosensitive channels dominate the minimal ion channel repertoire in prokaryotes

**DOI:** 10.64898/2026.03.26.714534

**Authors:** Cristóbal Uribe, Luciano Peña, Samuel Morales-Navarro, Sebastián E. Brauchi, Gonzalo Riadi, Juan C. Opazo, Wendy González

## Abstract

The eukaryotic genomes encode hundreds of proteins that function as ion channels and transporters. Essential for sustaining life, these proteins mediate the movement of inorganic ions (e.g., K^+^, Na^+^, Cl^−^, and Ca^2+^) across the plasma membrane according to their electrochemical gradients. In multicellular organisms, a diverse array of ion channels contributes to the maintenance of the resting membrane potential, the regulation of pH, osmolarity, and cell volume, and the control of secretion, electrical excitability, and synaptic activity, among many other fundamental physiological processes. Although independent evolutionary origins have been proposed for several ion channel families, their relative hierarchical importance for cellular viability remains poorly understood. To advance our knowledge of ion channel evolutionary history, we focused on determining the minimal combination of permeabilities that allows cellular viability. To this end, we conducted a survey of representative prokaryotes with small genomes across bacterial and archaeal phyla. By focusing on the smallest genomes, our approach enabled the identification of five ion channel architectures shared among prokaryotes. Among these, non-selective mechanosensitive channels (MscS and MscL) are the most abundant, followed by CLC-type channels, RCK-containing 2TM potassium channels, and proton channels of the MotA/TolQ/ExbB family. The conservation of the mechanosensitive protein architecture across archaeal and bacterial membranes suggests that the capacity to monitor physical membrane integrity predates the requirements for electrical communication.

## INTRODUCTION

Water, carbon, nitrogen, phosphorus, and essential ions -including potassium, calcium, protons, and chloride-, as well as electron acceptors such as oxygen or ferrous, together with appropriate temperature, are the essential elements that sustain life on earth. The regulated movement of ions across cellular membranes, both into and out of the cell, is an essential biological process that maintains cellular homeostasis and stability (Armstrong, 2003; Tosteson & Hoffman, 1960). This process involves both passive and active ion transport, mediated by specialized membrane proteins such as ion channels, transporters, and pumps (Gadsby, 2009). The relative impermeability of biological membranes to ions contrasts with the rapid permeability of water (Finkelstein, 1976). Essential for cell viability, ion transport alleviates the osmotic pressure generated by intracellular macromolecules, thereby protecting the structurally fragile cell membrane (Hille, 2018; Stigter & Hill, 1959). Ion selectivity, the preferential permeation of specific ions, enables efficient charge separation across the membrane (Hille, 2011). This electrical asymmetry is fundamental for establishing the resting membrane potential, which drives membrane transport, supports cellular metabolism, and underlies cellular excitability (Armstrong, 2015; Hille, 2011).

In multicellular organisms, membrane transport systems support both the electrical activity of excitable cells and the epithelial function (Hille, 2001). However, the ancient selection pressure that gave rise to ion channels as we know them today was likely different from the demands of electrical communication. It has been proposed that selective pathways for potassium, chloride, and protons constitute a minimal functional set required to sustain fundamental cellular functions, including osmotic balance, pH regulation, and the generation of electrochemical gradients that ultimately drive ATP synthesis (Armstrong, 2015). Compared with prokaryotic cells, eukaryotic cells are thought to have undergone multiple gene duplication events throughout their evolutionary history (Vosseberg et al., 2020). These events gave rise to an expanded repertoire of gene families involved in membrane permeability, including the H+ ATPase pumps and voltage-gated ion channels, enabling increasingly refined regulation of cellular and physiological functions (Gogarten et al., 1989; Makarova, 2005; Martins et al., 2013; Zakon, 2012).

The current availability of well-curated whole-genome sequences in public databases provides an unprecedented opportunity to annotate and investigate the diversity of ion channels. Over the past two decades, several high-throughput computational approaches have been developed to identify and annotate ion channels using genomic and transcriptomic datasets (Gao et al., 2016, 2019; Han et al., 2019; Lin & Ding, 2011; L.-X. Liu et al., 2006; Sankari & Manimegalai, 2017; Tiwari & Srivastava, 2015; Uribe et al., 2024; Zhao et al., 2017). However, despite these efforts, a comprehensive and systematic identification and comparative analysis of channels in prokaryotic organisms has not yet been carried out carefully (Kuo et al., 2005; Martinac et al., 2008).

Given the fundamental importance of maintaining ionic balance, we hypothesized that prokaryotic organisms with small genome sizes would harbor the “most economic” set of ion-permeating proteins that sustain the respective organisms. In principle, by analyzing the smallest genomes currently available in public databases within a phylogenetic framework, we can identify the minimal repertoire of ion channels and classify shared structural features underlying membrane transport. Here, we examined the ion transport systems in representative organisms from Archaea and Bacteria, beginning by identifying the species with the smallest genome in each phylum. Then, we sought to identify conserved functional and architectural elements across the selected organisms. To achieve this, we developed a bioinformatic pipeline for ion channel annotation that integrates sequence-based analyses with available three-dimensional structural information. Using this strategy, we identified five prevalent ion channel architectures among prokaryotes with minimal genomes. These include nonselective mechanosensitive channels MscS and MscL, CLC-type channels, RCK-containing 2TM potassium channels and proton channels ExbB from the MotA/TolQ/ExbB family.

## Materials and Methods

### Definition of a small genome and species sampling

Our sampling strategy included the species with the smallest genome size from each bacterial and archaeal phylum. Taxonomic classification and genome size data were obtained from the Genome Taxonomy Database (GTDB), release 226 (Parks et al., 2022). To define what constitutes a small genome, we adopted a statistical criterion by independently retrieving genome size data for available bacterial and archaeal species from the National Center for Biotechnology Information (NCBI) (Sayers et al., 2022). We then filtered the dataset for each domain to include only complete RefSeq genomes, meaning high-quality, well-annotated genomic sequences. For bacterial genomes, this filtering yielded 5,880 genomes. Based on this dataset, we defined a small genome as one in the lowest 5% of the genome-size distribution, corresponding to genomes smaller than 1.9 Mb. For archaeal genomes, we identified 374 complete genomes annotated in the RefSeq database. Using this subset, we defined small archaeal genomes as those smaller than 1.61 Mb. Based on these criteria, 23 bacterial and 6 archaeal genomes were identified as meeting the statistical threshold for classification as small genomes (Supplementary Table 1).

### Identifying ion channels in bacteria and archaea

After identifying the 29 species to study, we downloaded the protein-coding sequences (.faa files) from the NCBI database (Sayers et al., 2026). Once all the protein files for each species were obtained, we identified transmembrane proteins in the proteomes using the TMHMM v2.0, which identifies alpha-helical structures in protein sequences (Krogh et al., 2001). To do so, we applied a filter to retain all proteins with two or more transmembrane domains (Figure 1). Next, we annotated conserved domains for each transmembrane protein using the NCBI Batch Web CD-search Tool (Marchler-Bauer et al., 2011) with the CDD database version 3.21 (J. Wang et al., 2023), applying an E-value threshold of 0.01 to filter the results. We then identified ion channels using a precompiled file listing conserved ion-channel domains derived from CDD entries. This list was intersected with our NCBI Batch Web CD-search results using an *in-house* Perl Script. The output of these steps was a file listing putative ion channels for each species, which we manually curated to eliminate false positives (Supplementary Table S2). To verify the functionality of each candidate, we consulted reference databases such as NCBI and GeneCards (Stelzer et al., 2016).

**Figure 1.**
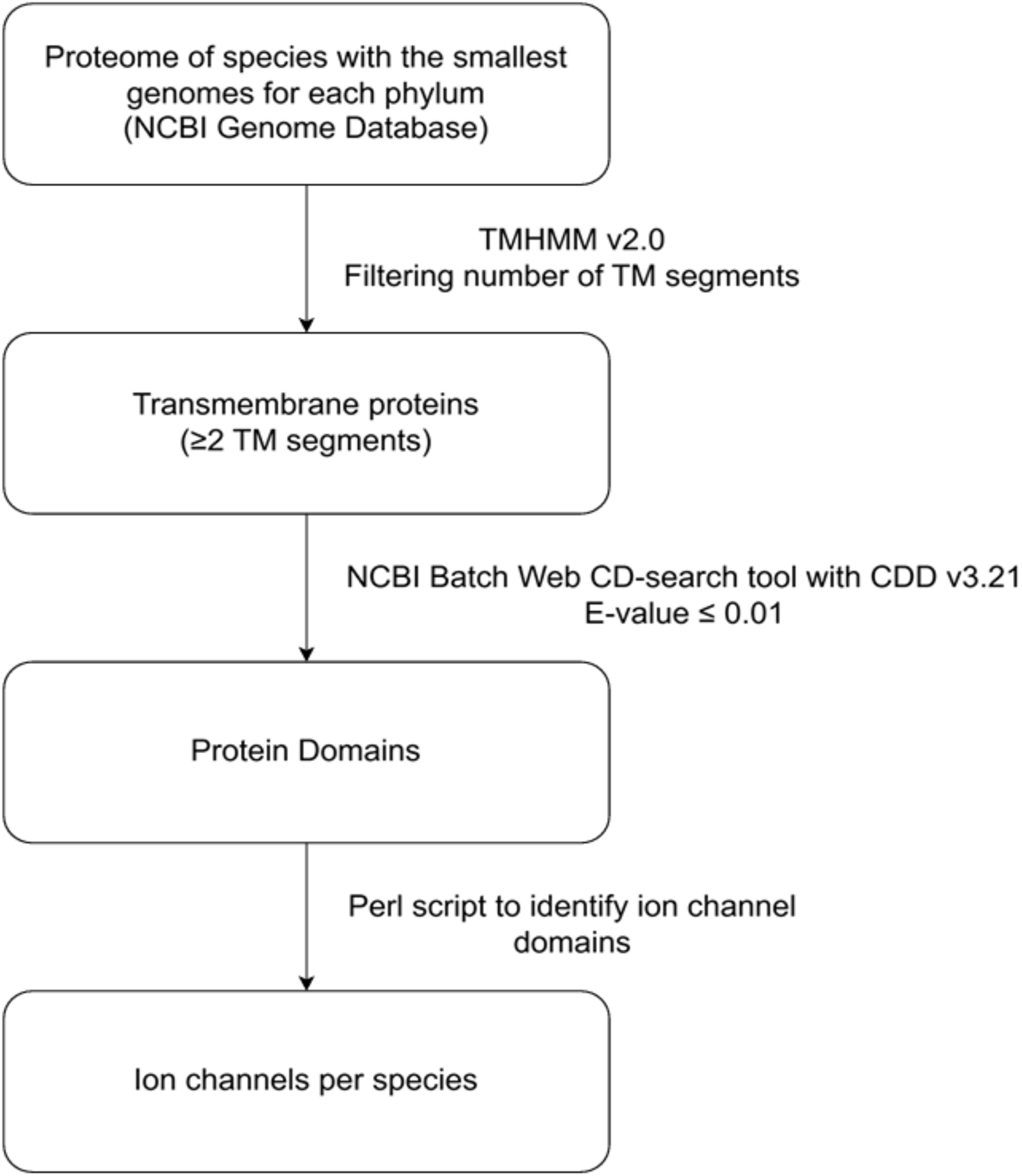
Workflow for identifying ion channels in small genomes. Proteomes from species with the smallest genomes *per* phylum were screened for transmembrane proteins (TMHMM v2.0) and annotated for conserved domains using the NCBI Batch Web CD-search tool with the Conserved Domain Database (CDD).

### Structural analysis

For each ion channel sequence identified using the previously described protocol, a protein BLAST (Camacho et al., 2009) search was performed against the Protein Data Bank (PDB) (Burley et al., 2023). For each query, the five highest-scoring segment pair (HSP) were selected, and the following filtering criteria were applied to choose a single hit: an E-value lower than 10^−6^, sequence identity greater than 30%, and alignment coverage greater than 40%. One representative structure *per* gene group was then manually chosen based on the frequency with which the same structure emerged as the top hit across group members (Supplementary Table S3). To assess structural conservation, amino acid sequences from each gene group were multiply aligned with their corresponding representative structures using Linux Version 7.526 (Multiple Alignment using Fast Fourier Transform) (Katoh, 2002). Positional conservation was quantified from the resulting alignment using a formula based on Shannon entropy: 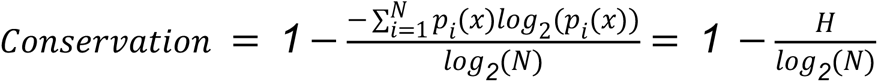 where *pi* represents the relative frequency of residue *i* at a position in the alignment, and *N* = 20, the number of standard amino acids. Conservation was defined as 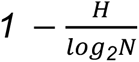, so a fully conserved position has a conservation value of 1, and a position with maximum variability has a conservation value of 0. Finally, normalized conservation values (ranging from 0 to 1) were mapped onto the selected three-dimensional structures (PDB entries shown in Figure 4) using a color gradient from red (low conservation) to blue (high conservation) in VMD (Humphrey et al., 1996). The same scale was applied to highlight conserved residues in the ClC and MthK selectivity filters, the MscL permeation pathway, and the pores of YnaI and ExbB (Figure 4, A to F, panel 4).

## Results

### Limited ion channel repertoires in small-genome prokaryotes

As shown in Figure 2, genome sizes across all examined species range from 0.15 Mb to 1.8 Mb, with corresponding protein-coding gene counts on the order of only a few hundred to ∼1,800. Despite this wide range in gene count, the number of identified ion channel genes remains low in all cases (Figure 2). Most of these minimal-genome organisms encode 0-4 ion channels (Figure 2). The highest value was estimated for the bacterial species *Thermosulfidibacter takaii* ABI70S6, which was 10 (Figure 2). In contrast, three bacterial species sampled lacked ion channels in their protein-coding gene repertoires (Figure 2). Thus, ion channels constitute a small fraction of the protein-coding gene repertoire in the sampled species, accounting for less than 1% of all protein-coding genes (often just 0.1–0.5%, when present at all). Thus, our results reveal that prokaryotic organisms with the smallest genome sizes possess a limited repertoire of ion channels or use entirely unreported ones.

**Figure 2.**
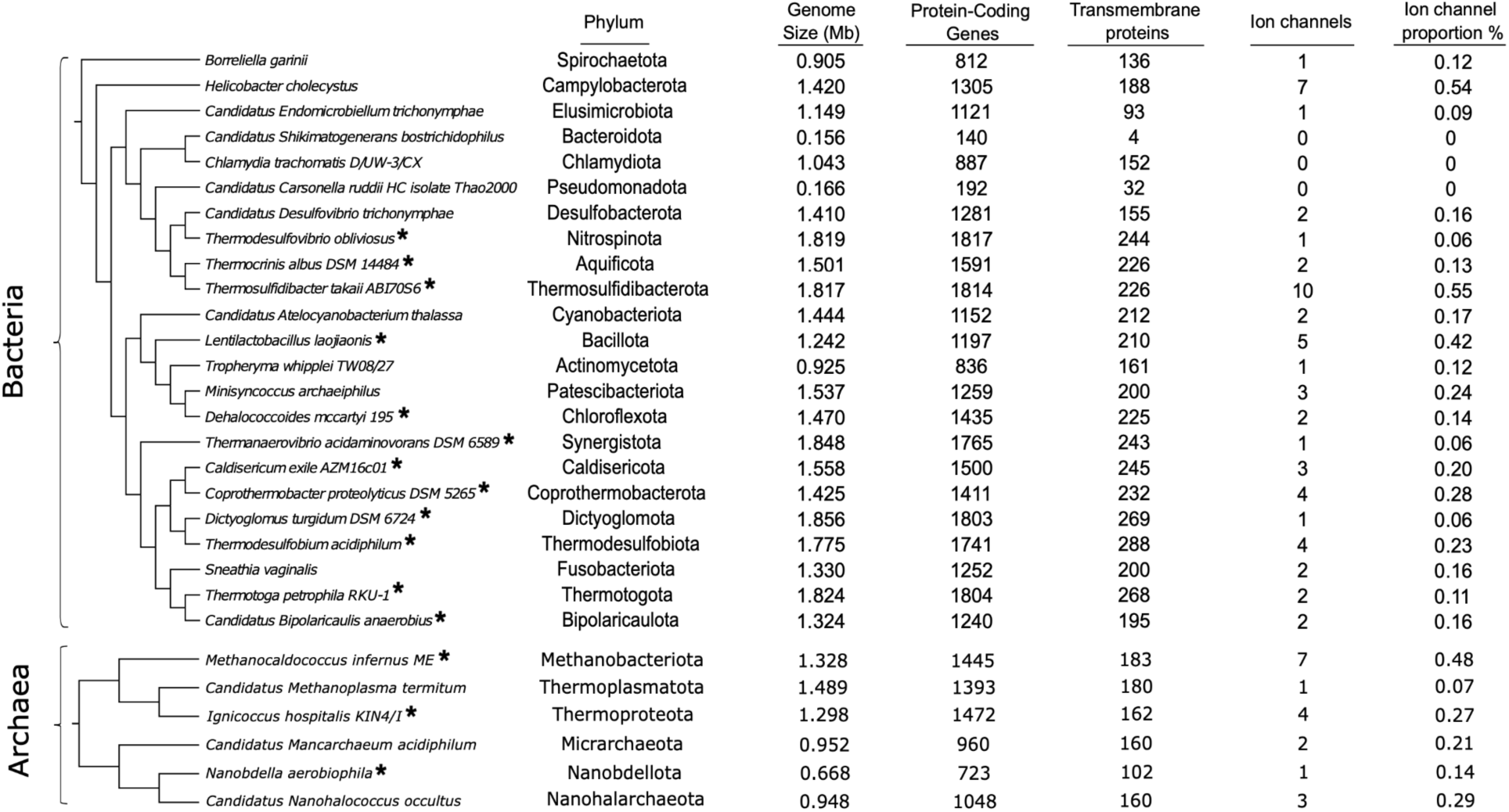
Phylogenetic trees of the bacterial and archaeal species included in our study. For each species, the phylum, genome size, number of protein-coding genes, number of transmembrane proteins, number of identified ion channels, and the proportion of ion channels relative to the total number of protein-coding genes are shown. Asterisks (*) denote species classified as free-living.

### Free-living vs. non-free-living bacteria differ in ion channel gene counts but not in their proportions

Within the bacterial group, we observed that although the average proportion of ion channels is not different between free-living (0.19%) and non-free-living (0.19%) species (z-test for the difference between proportions, p-value = 0.485, Supplementary Table S4, the total number of channels is indeed different (free-living species, n = 12, total number of ion channels = 37; non-free-living species, n = 11, total number of ion channels = 19). The lack of difference in the proportion is likely due to the fact that free-living bacteria have, on average, more protein-coding genes (n = 12, with an average of 1593.17 protein-coding genes) compared to non-free-living species (n = 11, with an average of 930.64 protein-coding genes) (Welch t-student test p-value = 0.0003, Supplementary Table S4). In our dataset, all three bacterial species that we estimate lack ion channels in their protein-coding gene repertoires are non-free-living (Figure 2). Two of them have the smallest genome sizes reported in our sampling, *Candidatus Carsonella ruddii* and *Candidatus Shikimatogenerans bostrichidophilus* (Figure 2). For species with 1, 2, or 3 ion channels encoded in their genomes, half are free-living (Figure 2). For species that have 4 or more ion channels, most are free-living (Figure 2). In summary, bacterial species with extreme genome reduction, often non-free-living, may have eliminated canonical ion channels from their genomes—an outcome generally considered unlikely—or may instead rely on as yet unreported or highly divergent ion channels. For those with an intermediate number of ion channels (1 to 3), there is no bias in their lifestyle. In contrast, for those with the highest number (4 to 10), there is indeed a bias, as most are free-living species.

### Free-living archaea tend to encode more ion channels despite a similar number of protein-coding genes

The small-genome archaea follow patterns similar to those of their bacterial counterparts, with uniformly low ion channel gene content (Figure 2). Although the uneven sample sizes have prevented seeing a statistically significant difference between the proportion of ion channels between free-living (0.33%) and non-free-living (0.18%) species (z-test for the difference between proportions, p-value = 0.23, Supplementary Table S4), the first group nearly doubles the second one in proportion of ion channels. Interestingly, and contrary to the pattern found in bacteria, the smallest genomes of archaea do not have a contrasting number of protein-coding genes (n = 3, with an average of 1,213.33 protein-coding genes for free-living species and n = 3, with an average of 1,133.67 protein-coding genes for non-free-living species, Welch t-student test p-value = 0.79, Supplementary Table S4). All six archaeal genomes analyzed are ∼1.5 Mb or less (the smallest being ∼0.67 Mb) and encode only 1–7 ion channel genes each (Figure 2). The most reduced archaeal genome in our dataset (e.g., *Nanobdella aerobiophila*, ∼0.67 Mb, 723 protein-coding genes) retains one ion channel gene (Figure 2). On the other hand, the hyperthermophilic methanogen *Methanocaldococcus infernus* (1.33 Mb, 1,445 genes) encodes 7 ion channels, the largest number among the examined archaean species (approximately 0.48% of its genes). In our dataset, species with the fewest ion channels, ranging from 1 to 3, are predominantly non-free-living (Figure 2). In contrast, species with the largest number of ion channels, ranging from 4 to 7, are mostly free-living species (Figure 2). Thus, the smallest archaeal genomes tend to have a lower proportion of ion channels, with higher proportions in free-living species.

### Mechanosensitive channels dominate ion channel repertoires in small-genome prokaryotes

After classifying ion channels in bacterial and archaeal species with small genomes, we found that mechanosensitive channels are the most represented type (Figure 3). Notably, among the eight species that encode a single ion channel, seven possess a mechanosensitive channel (Figure 3). The exception is *Thermanaerovibrio acidaminovorans* DSM 6589, which harbors a single potassium channel. Among species with two ion channels, we found a similar pattern: in all but one case, one channel is mechanosensitive (Figure 3). The second channel is typically either a potassium or proton channel in roughly equal proportions, with one species encoding a chloride channel as the second type. Interestingly, *Candidatus Bipolaricaulis anaerobius* is the only species in which both ion channels are mechanosensitive. In species encoding three ion channels, two (one bacterium and one archaeon) have exclusively mechanosensitive channels, while the third species encodes two mechanosensitive channels and one chloride channel (Figure 3). These findings suggest that mechanosensitive channels are the most prevalent type among prokaryotes with small genomes, a trend particularly pronounced in species with a limited repertoire of ion channels.

**Figure 3.**
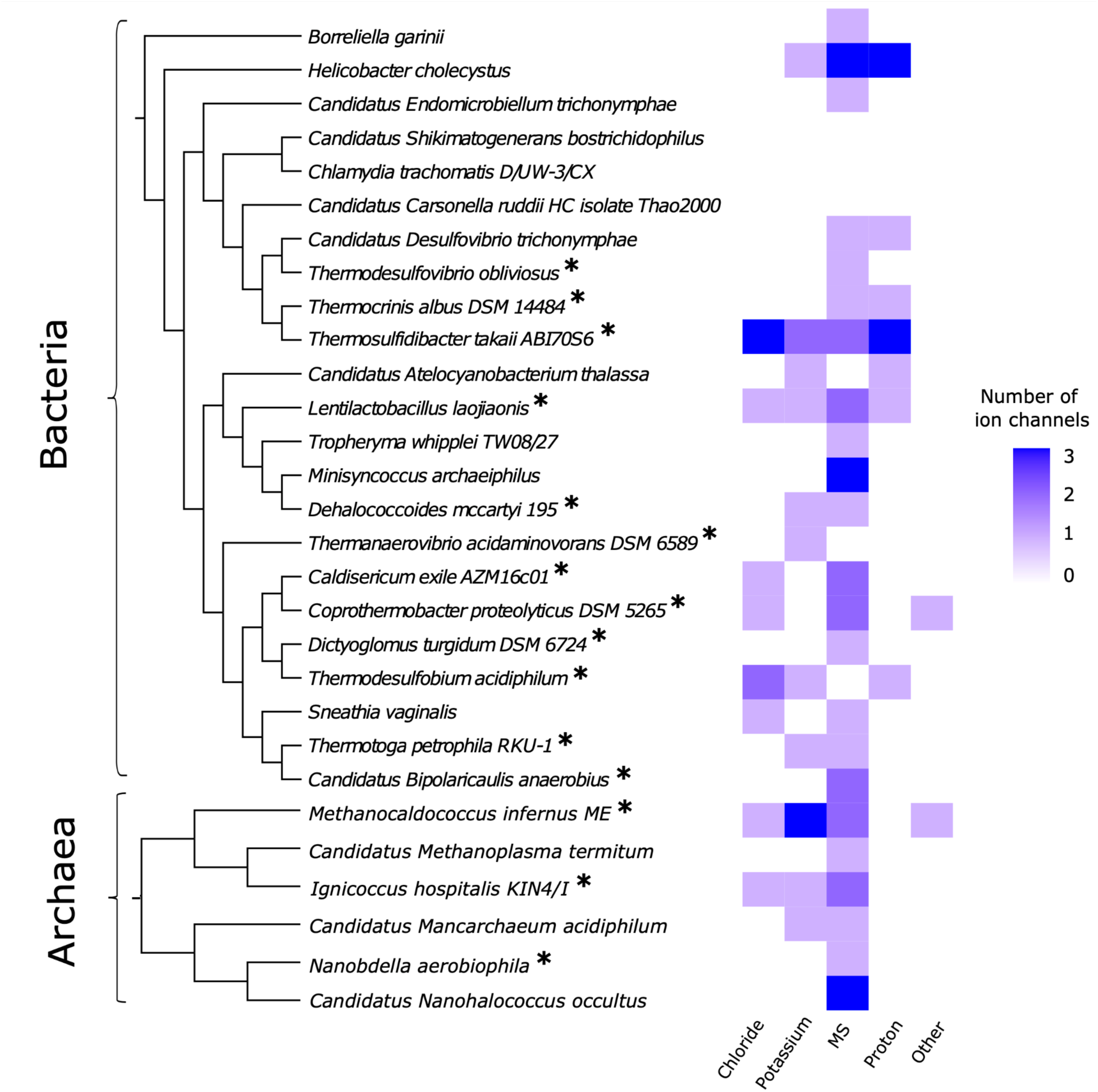
Heatmap displaying the number and types of ion channels identified in each species included in our study. MS, Mechanosensitive channels. Asterisks (*) denote species classified as free-living (Supplementary Table 2).

### Structural conservation of the ion channel set of minimal genomes

By analyzing multiple sequence alignments of the ion channels against Protein Data Bank targets (Supplementary Figure 1), we determined their conservation across the five prevalent architectures found in our study (Figure 4).

**Figure 4.**
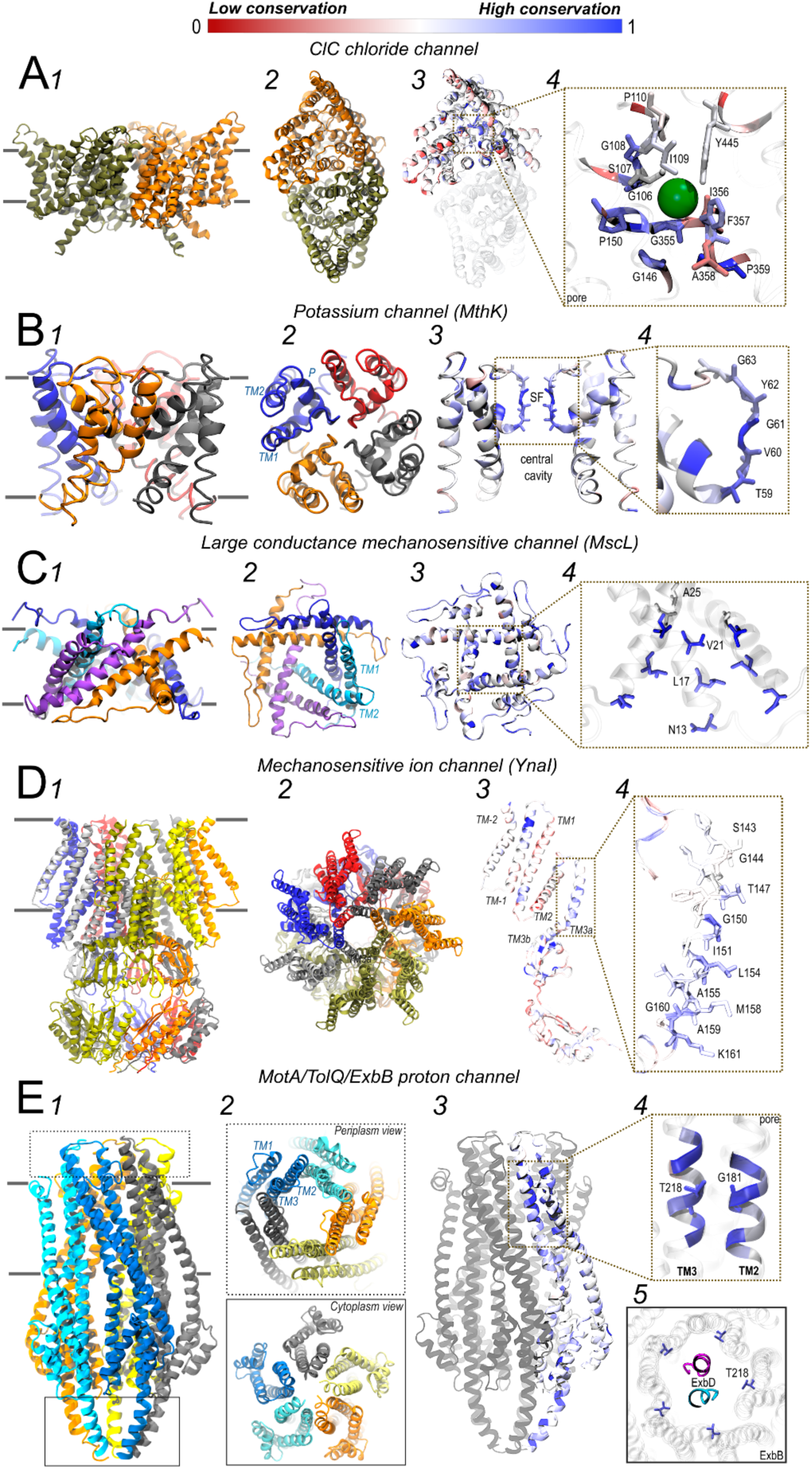
Structures of the five prevalent ion-channel architectures and their conservation patterns. A) Chloride Channel (ClC) (PDB code: 1KPL), B) Potassium channel, pore domain MthK channel (PDB code: 1LNQ), C) Large conductance mechanosensitive channel SaMscL(CΔ26) (PDB code: 3HZQ), D) Low conductance mechanosensitive channel YnaI (PDB code: 6URT) and E) Coupled proton channel, ExbB/ExbD (PDB code: 7AJQ). Subunits are represented by different colors in panels 1 and 2 of each figure. In panel 1, horizontal gray lines represent the boundaries of the helices to the membrane. Panel 2 shows the channel from a top perspective, except for ExbB. Panel 3 shows the conservation (see methods) for each channel. Panel 4 zooms in on the selectivity filter region for ClC (Figure A) and MthK (Figure B), as well as the pore region for MscL (Figure C), YnaI (Figure D), and ExbB (Figure E). Panel 5 shows the ExbB/ExbD complex. Conservation is shown in a spectrum from blue (highest conservation) through white (medium conservation) to red (lowest conservation).

CLC-type channels align with the ClC chloride channel from *S. typhimurium* (PDB ID: 1KPL) and the *Synechocystis sp.* ClC channel (PDB ID: 3ND0). These channels facilitate the passive and selective transport of anions, primarily Cl⁻, across the lipid bilayer. The crystallographic structure of the ClC channel reveals a homodimeric organization (Figure 4A, panels 1 and 2), where each subunit contains an independent ion conduction pore. Each monomer comprises 18 transmembrane helices (TMs) arranged in an antiparallel topology, generating internal pseudo-symmetry (Jayaram et al., 2011). As shown in Figure 4A, structural conservation was primarily observed in the central regions of the transmembrane (TM) segments, with particularly high conservation in the anion selectivity filter (SF) region (Figure 4, panel 4). This SF includes the motifs GSGIP (residues 106–110), GREGP (146–150), GIFAP (355–359), and the Y445 residue. Within these motifs, residues such as G106, G108, G146, P150, G355, F357, and P359 exhibit conservation scores above 0.8.

The potassium channel group aligns with the MthK channel from *Methanobacterium thermoautotrophicum* (PDB ID: 1LNQ). This channel assembles as a homotetramer, with each subunit contributing a pore-forming domain comprising two TMs helices and two intracellular RCK (Regulator of Conductance for K⁺) domains (Jiang et al., 2002). The pore domain adopts a canonical architecture that is highly conserved across potassium channels (Figure 4B, panels 1 and 2). A hallmark of this domain is the SF, formed by a loop between TM1 and the P domain (pore helix) of each subunit (Figure 4B, panels 2 and 3). This loop contains the signature TVGYG motif (Figure 4B, panel 4), which is highly conserved among K⁺ channels and essential for selective ion conduction. This motif shows calculated conservation values of 0.90, 0.84, 1, 0.80, and 0.75, respectively. The backbone carbonyls and side chain oxygen atoms of this motif are spatially arranged to coordinate dehydrated K⁺ ions via a precise geometry that mimics their hydration shell, enabling efficient conduction while excluding smaller ions such as Na⁺.

One of the identified mechanosensitive channel types corresponds to the large-conductance channel (MscL). The amino acid sequences of the genes analyzed showed more than 33% identity to the reference structure. As a representative structure, the MscL channel of *Staphylococcus aureus* was selected (PDB ID: 3HZQ), whose crystallographic structure corresponds to a truncated variant representing an expanded intermediate state of the channel (SaMscL(CΔ26)). However, independent of this conformation, the MscL channel adopts a tetrameric organization, in which each subunit presents two TMs separated by a loop (Figure 4C, panels 1 and 2). On the other hand, despite differing in the number of subunits from the *Mycobacterium tuberculosis* pentameric channel (MtMscL), also identified in this study in the query of the sequences against PDB (PDB ID: 2OAR), the relative arrangement of transmembrane segments and pore topology is conserved (Supplementary Figure 2). The pore domain in MscL channels is highly conserved since it constitutes the functional backbone of the channel (Figure 4C, panels 3 and 4). The narrowest point of the pore in SaMscL(CΔ26) is located near residue V21, flanked by L17 and A25, which are also conserved among the analyzed sequences and help delineate the channel permeability profile. Unlike potassium channels, MscL lacks a classical selectivity filter. Its opening generates a large-diameter pore, whose maximum width is approximately around the N13 residue. The estimated evolutionary conservation for key residues N13, L17, V21, and A25 was 0.86, 0.80, 1.0, and 0.48, respectively, highlighting the functional importance of this region in pore architecture and function (Z. Liu et al., 2009; Steinbacher et al., 2007).

Other identified mechanosensitive channels aligned best with mechanosensitive small-conductance channel (MscS)-like structures. The results indicated that the *E. coli* YnaI channel (PDB ID: 6URT) is the best structural representative. This channel is a paralog of the MscS channel from *E. coli*, sharing a membrane voltage-dependent activation mechanism, although it presents a more complex transmembrane architecture (Steinbacher et al., 2007). The structure of YnaI resolved corresponds to a nonconducting state and reveals a heptameric assembly (Figure 4D, panels 1 and 2), in which each subunit contributes five transmembrane helices, numbered from TM-2 to TM3 (Figure 4D, panel 3). Among these, helix TM3a, composed of residues L142-A159, lines the central pore and constitutes the main structural component of the ionic conduction pathway. This helix forms a heptameric bundle together with its adjacent subunit counterparts (Figure 4D, panel 2), generating a predominantly hydrophobic environment in the pore lumen. Within TM3a, residues L154 and M158 form two successive hydrophobic layers that define the channel bottleneck (Figure 4D, zoom in panel 3), analogous to residues L105 and L109 of the *E. coli* MscS channel (Supplementary Figure 3). Additionally, residue G160, immediately downstream of TM3a, introduces a structural break in the helix, allowing the side chain of K161 to orient toward the center of the pore, generating a third constriction with a possible role in the cation selectivity of the channel (Hu et al., 2021). This set of residues, particularly those oriented toward the center of the pore, is moderately conserved across the analyzed genes of the mechanosensitive channel family. The conservation values obtained were: T147 (0.71), G150 (0.86), I151 (0.71), L154 (0.80), M158 (0.71), A159 (0.80), K161 (0.76). These values suggest a functional convergence in the formation of the mechanosensitive channel pore (Figure 4D, panel 3).

Finally, for the genes identified within the family of proton channels (MotA/TolQ/ExbB), the *Serratia marcescens* ExbB channel structure (PDB ID: 7AJQ) was selected as a representative. ExbB is part of the Ton complex, a transmembrane machinery that transduces proton motive force (pmf) energy from the inner membrane to specific outer membrane transporters. This complex is composed of ExbB, ExbD, and TonB (or HasB in *S. marcescens*) (Biou et al., 2022). The structure of ExbB corresponds to a pentameric assembly (Figure 4E, panels 1 and 2), in which each subunit contributes with three highly conserved TMs (Figure 4E, panel 2), which together delimit a central pore of apolar character. This organization agrees with that observed in *E. coli* ExbB/ExbD complex (e.g., PDB ID: 5SV0) (Supplementary Figure 4), suggesting evolutionary structural conservation of the ExbB assembly. This conservation is also reflected in the homology between ExbB/ExbD and the stator elements (MotA/MotB) of the bacterial rotary motor, which share a common mechanism for coupling the proton motive force (pmf) to mechanical work (Lai et al., 2020). In this context, two residues have been described that could play key roles in the architecture of the ExbB/ExbD complex and in modulating proton transduction. In particular, residues G181 and T218, located in TM2 and TM3 of ExbB, respectively (Figure 4E panel 4). The residue T218 orients its polar side chain toward the pore lumen, suggesting a possible involvement in stabilizing the interaction with ExbD helices (Figure 4, panel 5). High evolutionary conservation was observed for both residues, with values of 1.0 and 0.85 for G181 and T218, respectively. Although ExbB does not function as a channel on its own, it is an essential component of the energy-generating and proton-conducting system ExbB/ExbD/TonB. Electrophysiological and mutational studies in *E. coli* have shown that the ExbB/ ExbD complex exhibits pH-dependent cation-selective conducting activity and that the hydrophobic pore of ExbB is occupied by the transmembrane helices of ExbD in functional states (Figure 4, panel 5), suggesting a gating mechanism or conformational control coupled to pmf usage (Celia et al., 2016; Maki-Yonekura et al., 2018).

## Discussion

A growing body of research in organisms with minimal genomes has enabled the identification and classification of genes that are essential for life (Akerley et al., 2002; Baba et al., 2006; De Berardinis et al., 2008; French et al., 2008; Giaever et al., 2002; Glass et al., 2006; Ji et al., 2001; Kobayashi et al., 2003; Langridge et al., 2009; Liberati et al., 2006; Salama et al., 2004). Within this repertoire, genes involved in cellular membrane biogenesis are unquestionably critical for viability, as they enable the segregation of protoplasm from the extracellular milieu and promote cellular specialization through compartmentalization of biological processes and regulated solute fluxes that maintain cellular homeostasis, with ion channels traditionally viewed as central mediators of these functions.

### The Paradox of Ion Channel Dispensability and Conditional Essentiality

In this study, we applied an integrated phylogenetic and structural framework to define the minimal repertoire of ion channels required for prokaryotic life. Our findings indicate that this repertoire comprises five molecular architectures—MscS, MscL, ClC, potassium channels (2TM-RCK), and ExbB proton channel—present even in the most genomically reduced organisms.

Notably, a striking finding from our analysis is the very low proportion of ion channel genes, which, on average, account for only 0.19% of annotated coding sequences in the minimal-genome species included in our study. This estimated proportion contrasts sharply with the approximately 0.96% reported for a representative group of mammals (Uribe et al., 2024). In some cases, ion channel genes appear to be entirely absent, as observed in obligate symbionts such as *Candidatus carsonella ruddii*. Consistent with this pattern, parasitic or symbiotic bacteria generally tend to lack K⁺ channels (Kuo et al., 2005). These observations indicate that bacterial species with extreme genome reduction, often non-free-living, may have either largely eliminated canonical ion channels from their genomes—an outcome generally considered unlikely—or instead rely on as yet unreported or highly divergent channel-like mechanisms.

Importantly, the identified ion channel common set does not appear to constitute a critical component of electrical excitability or complex signalling mechanisms in these simple organisms, unlike in metazoans. Rather, its likely functions as an optimized kit that preserves cellular integrity and meets fundamental energetic requirements.

Together, these results challenge the hypothesis that ion channels are conditionally essential, particularly for free-living organisms in fluctuating environments (Breuer et al., 2019; Glass et al., 2006). In this context, ion channels may become dispensable in highly stable environments, such as the cytoplasm of host cells or under controlled laboratory conditions, as exemplified by the synthetic minimal organism JCVI-syn3.0 (Breuer et al., 2019), where ionic requirements may be satisfied through passive solute fluxes or effectively delegated to the host cell’s transport machinery.

The conspicuous absence of ion channels in *C. ruddii* further suggests that, under conditions of maximal genomic reduction, alterations in membrane lipid composition may become functionally critical, serving as a sufficient diffusion barrier, as proposed for membrane transport in primitive cells (Mansy, 2010). Under such circumstances, metabolite transport and the fulfilment of energetic demands would depend primarily on a limited repertoire of active transporters rather than on a minimal ensemble of channels dedicated to maintaining the membrane potential (Justice et al., 2024; McCutcheon & Moran, 2012; Nakabachi et al., 2006). Accordingly, the apparent dispensability observed in our study likely reflects an evolutionary adaptation to specific environmental contexts, in which the energetic cost of maintaining osmotic gradients through selective channels may outweigh cellular benefits, thereby favouring alternative strategies such as active transport systems or passive diffusion across an adapted lipid bilayer. Besides, our pipeline was not designed to predict small membrane proteins containing a single transmembrane segment (Yadavalli & Yuan, 2022), which could also contribute to ionic balance in these minimal organisms. Although many bacterial pore-forming proteins (PFPs) of a single transmembrane segment are secreted as soluble monomers that bind to host cell membranes, oligomerize, and form transmembrane pore complexes that disrupt plasma membrane integrity (Chatterjee et al., 2025), we cannot exclude the possibility that related proteins may also form ionic pores in minimal-genome bacteria. Our methodology was also not designed to identify proteins with β-sheet–based structures, which may also have influenced our results. Indeed, membrane proteins adopting β-barrel topologies have been described in the outer membranes of Gram-negative bacteria, as well as in plastids and mitochondria (Duy et al., 2007; Paschen et al., 2003). These proteins perform a wide range of biological functions, including roles as porins, transporters, enzymes, virulence factors, and receptors (Fairman et al., 2011), raising the possibility that they may contribute to ionic balance.

### Mechanosensitivity as an Ancestral Safety Valve

The higher abundance of mechanosensitive channels (MscL and MscS) observed in our survey (Figure 3) underscores that protection against physical stress is likely a primary selective pressure, rather than chemical signaling. In earlier analyses of essential genes in minimal bacteria, such as *Mycoplasma genitalium*, other channel types, including Trk potassium transport systems, were identified as essential (Glass et al., 2006), however, mechanosensitive channels were not explicitly discussed in those studies. Unlike voltage-or ligand-activated channels, which participate in highly sophisticated transduction cascades, mechanosensitive channels respond directly to a more general signal—lipid bilayer tension (Martinac et al., 2008). From a metabolic perspective, MscL represents an optimal solution for minimal genomes, as its hallmark characteristics of high conductance and low selectivity (Kloda & Martinac, 2002) enable it to function as a pressure-relief valve during abrupt changes in extracellular solute concentration, thereby preventing osmotic lysis without ATP expenditure or reliance on complex molecular networks. The conservation of this protein architecture across archaeal and bacterial membranes (Figures 4C and 4D) suggests that the capacity to monitor physical membrane integrity predates the requirements for electrical communication. While the loss of selective channels can be metabolically compensated, the absence of mechanosensitive elements during osmotic fluctuations is typically lethal, thereby justifying their evolutionary prioritization.

### Evolutionary Convergence of Ion Permeation Mechanisms in Minimal Channel Systems

The structural analysis presented herein demonstrates that, despite genomic reduction eliminating redundancy and accessory subunits, the chemical principles governing ionic permeability are remarkably conserved (Doyle et al., 1998). A strict evolutionary convergence appears to define channel selectivity and gating mechanisms (Sukharev & Corey, 2004). Regarding anionic selectivity, ClC channels demonstrate clear conservation of the GSGIP (106–110), GREGP (146–150), and GIFAP motifs (355–359) (Figure 4A), specifically residues G106 and G108 from the first motif, G146 and P150 (from the second) and G355, F357 (from the third), confirming persistence of an ancestral anionic coordination mechanism through these motifs. They are placed at the N termini of α-helices, pointing towards the binding site (Figure 4A, panel 4). This arrangement of helices is expected to create an electrostatically favourable environment for anion binding, mediated by interactions with the amide group of F357, thereby reducing the energetic burden of introducing stronger positive charges within the pore (Dutzler et al., 2002).

Regarding dehydration geometry, MthK-like potassium channels conserve the canonical TVGYG motif (Figure 4B), demonstrating that coordination of fully dehydrated K⁺ ions is a universal requirement to avoid loss of selectivity against competing ions such as Na⁺ (Jiang et al., 2002). Within this conserved motif, the third residue, a glycine, is fully conserved in all analyzed sequences (Supplementary Figure 1). This residue contributes to the formation of sites S1 and S2 of the selectivity filter, which are essential for K⁺ coordination. Loss of K⁺ coordination at these two sites favors Na⁺ permeation (Andrini et al., 2024).

The high conservation of L17 and V21 residues in MscL channels, at the hydrophobic pore constriction (Figure 4C) reveals a gating mechanism characterized by physicochemical simplicity through nonpolar amino acids, rendering it readily encodable within minimal genomes (Sukharev & Corey, 2004). In contrast, the homolog residues at MscS channels, L154 and M158 (Figure 4D), are less conserved. Finally, in the ExbB proton channel from the MotA/TolQ/ExbB family, the presence of a small and a specific polar amino acid (G181 and T218, respectively) within an otherwise nonpolar transmembrane region (Figure 4E) suggests that energetic coupling requires precise electrostatic motifs facilitating the proton flux essential for active transport of molecules. In particular, T218 faces Asp25 from ExbD. Asp25 connects the cytoplasm to the periplasm and has been proposed as an essential residue for proton transfer through these channels (Biou et al., 2022) (Biou et al., 2022). G181 is a threonine in the ExbB proton channel from *E. coli*, and mutagenesis studies on this residue have revealed its importance for harnessing the pmf (Celia et al., 2016).

### The Cost of Complexity

In the genomically reduced organisms analyzed in this study, the low abundance of ion channels may be tolerated—or even compensated by the host or environmental conditions. In contrast, in larger and more complex multicellular organisms, mutations in homologs of the minimal channel repertoire identified here can lead to severe and often lethal pathologies, including cystic fibrosis and cardiac arrhythmias (Hatta et al., 2002). This striking contrast suggests a functional divergence in which, while in minimal genomes ion channels primarily serve as robust mechanisms for cell survival, in complex organisms they have been co-opted for higher-order physiological signalling, revealing an increased vulnerability associated with genomic and functional complexity (Gibson et al., 2010). This notion is supported by the observation that lethality in humans typically arises not from the metabolic failure of individual cells but from the collapse of organismal communication networks (Sukharev & Corey, 2004; Zaydman et al., 2012). A pertinent example is the mutation of the sodium channel Nav1.5 (long QT syndrome), which does not directly cause cardiomyocyte death but instead induces fatal desynchronization of cardiac tissue (Arkhipov, 2025; Q. Wang et al., 1995). An equivalent bacterial defect would most likely manifest as reduced growth rate rather than lethality, thereby avoiding lethal consequences. Collectively, these arguments support the idea that the transition from unicellularity to multicellularity transformed ion channels from safety valves into components of systemic communication networks, helping explain why minimal genomes can lack elements that are indispensable for human survival (Liebeskind et al., 2015; Martinac et al., 2008).

JCVI-syn3.0 represents the first self-replicating organism with a minimized synthetic genome (473 genes, 531 kbp), developed by the J. Craig Venter Institute to identify essential genes and establish a platform for synthetic biology (Hutchison et al., 2016). In essence, it functions as a genome-wide knockout of non-essential genetic components. Notably, this organism lacks most members of the Kv, Nav, and Cav channel families, highlighting their dispensability for basic cellular viability. Subsequent work leading to the construction of JCVI-syn3A, achieved through the reintroduction of 19 genes into syn3.0 restore normal cellular morphology (Pelletier et al., 2021). Remarkably, none of the introduced genes encoded ion channels. Together, these synthetic genomics experiments demonstrate that fundamental organismal viability does not require ion flux mediated by canonical ion channels.

### The Role of Lipid Membranes in Minimal Organisms

The scarcity of ion channels in the organisms examined in our study raises a fundamental biophysical question: how is membrane potential maintained in the absence of protein mediated regulation of membrane permeability? Recent studies of the synthetsufficient-syn3A (Gan et al., 2025; Justice et al., 2024) demonstrate that a membrane composed solely of phosphatidylcholine and cholesterol can sustain cellular viability, provided that passive permeability remains sufficiently to preserve osmotic balance (Justice et al., 2024). These findings suggest a potential inverse coevolutionary relationship between the complexity of the ion channel repertoire and lipid membrane permeability. In the absence of an extensive network of channels and pumps regulating ionic flux, minimal-genome organisms may rely on tightly regulated lipid composition that minimizes passive permeability, effectively transforming the membrane into an active insulating barrier (Heimburg, 2010; Shinoda, 2016), a role that in more complex organisms is achieved through ion channels and ATP-dependent pumps. Such biophysical compensation would allow minimal cells to maintain essential electrochemical gradients despite their highly reduced repertoire of ion channels.

### Conclusions and Future Perspectives

In summary, our phylogenetic and structural analyses define a minimal ion channel repertoire dominated by the physical stress response (mechanosensitivity) and energetic coupling (ExbB/ExbD complexes). This repertoire does not represent a simplified version of the eukaryotic ion channel interactome, but instead reflects a fundamentally distinct adaptation to environmental constraints. The lethality of channelopathies in humans—including cystic fibrosis (chloride channel CFTR), long QT syndrome and arrhythmias (KCNQ1 and HERG potassium channels; Nav1.5 sodium channels), and episodic ataxias (Kv1.1 or Cav2.1) (Arkhipov, 2025; Hatta et al., 2002)—contrasts sharply with the apparent dispensability of these channels in minimal or synthetic cells, revealing that evolution repurposed the ancestral cellular survival machinery for intercellular communication functions in multicellular organisms. This reinforces the concept that electrical excitability represents a late evolutionary adaptation, fundamentally dispensable for the thermodynamic definition of cellular life.

Future experimental validation of these hypotheses will require approaches that extend beyond comparative genomics. In particular, synthetic models should incorporate not only the minimal channel repertoire but also native lipid composition characteristic of extremophile and symbiotic membranes. Understanding how lipid remodelling compensates for the scarcity of protein-based transport systems may open new avenues for rational nanoreactor design and contribute substantially to our understanding of the minimal physicochemical requirements underlying the origin and persistence of cellular life.

## Declaration of generative AI and AI-assisted technologies in the manuscript preparation process

During the preparation of this work the author(s) used ChatGPT (OpenAI) in order to improve grammar and language clarity. After using this tool, the author(s) reviewed and edited the content as needed and take(s) full responsibility for the content of the published article.

## Acknowledgments

The authors acknowledge the National Laboratory for High Performance Computing (Chile), Fondecyt Project 1230446 (W.G.), Fondecyt Project 1231357 (G.R.), Fondecyt Project 1241753 y PEW Innovation Fund (S.B), Fondecyt Project 1250688 (J.O.). C.U. and L.P acknowledge the ANID National Doctorate Scholarships 21241552 and 21231159 respectively.

## Supplementary figures

**Fig. 1.**
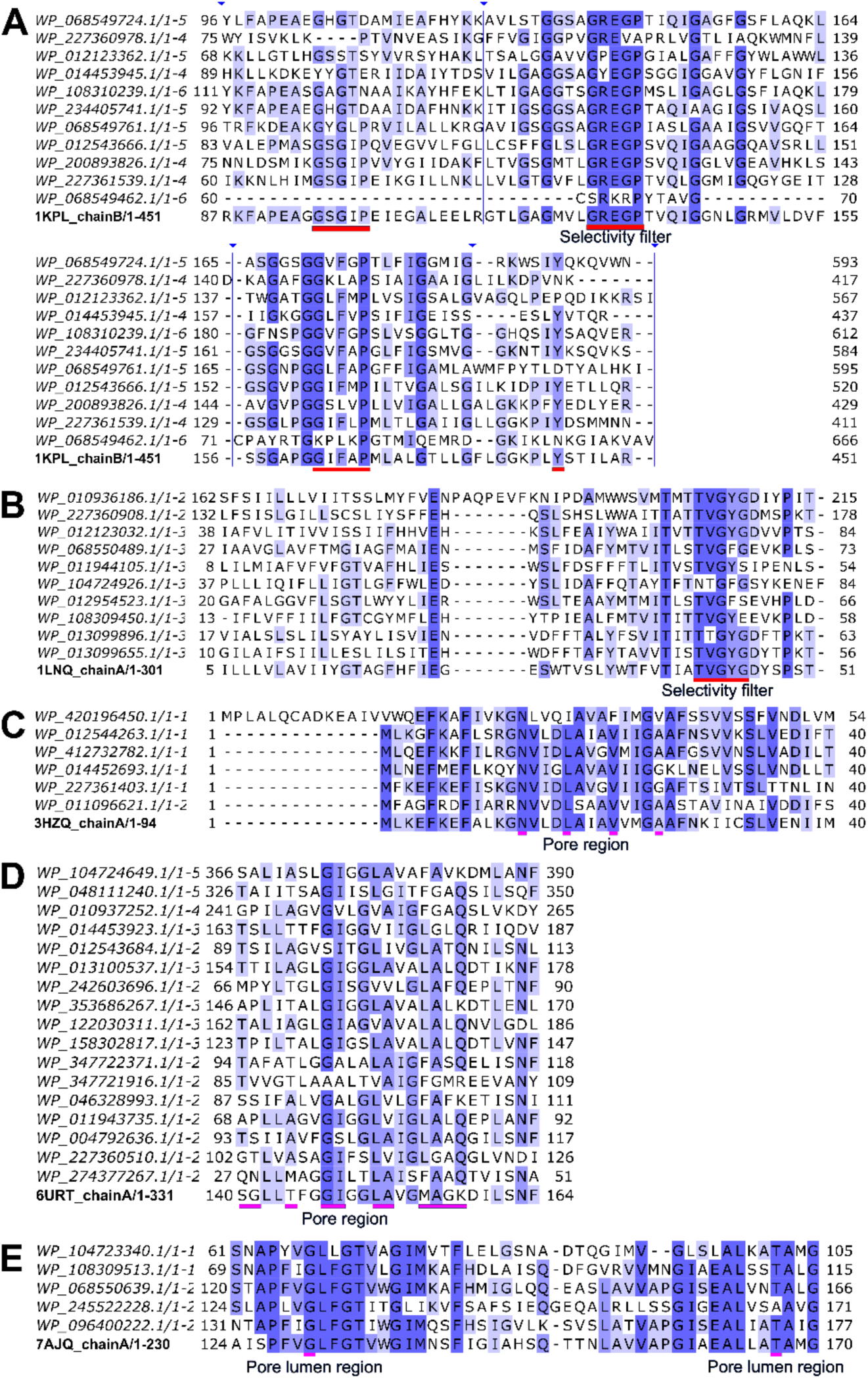
Multiple Sequences Alignment of ion channel protein sequences of five families identified in minimal genome organisms and representative structure sequence. A) Chloride Channel (ClC) (PDB code: 1KPL), B) Potassium channel, pore domain MthK channel (PDB code: 1LNQ), C) Large conductance mechanosensitive channel SaMscL(CΔ26) (PDB code: 3HZQ), D) Low conductance mechanosensitive channel YnaI (PDB code: 6URT) and E) Coupled proton channel, ExbB/ExbD (PDB code: 7AJQ). The alignment coloring is shown according to the percentage of identity; it does not represent conservation, as calculated and shown in Figure 4 (see Methods). The structural regions highlighted in our analyses are marked with a red box in A and B (selectivity filters), and pink box in C, D and E (pore regions). The thin blue vertical lines in A are segments of the sequences hidden by visual motifs.

**Fig. 2.**
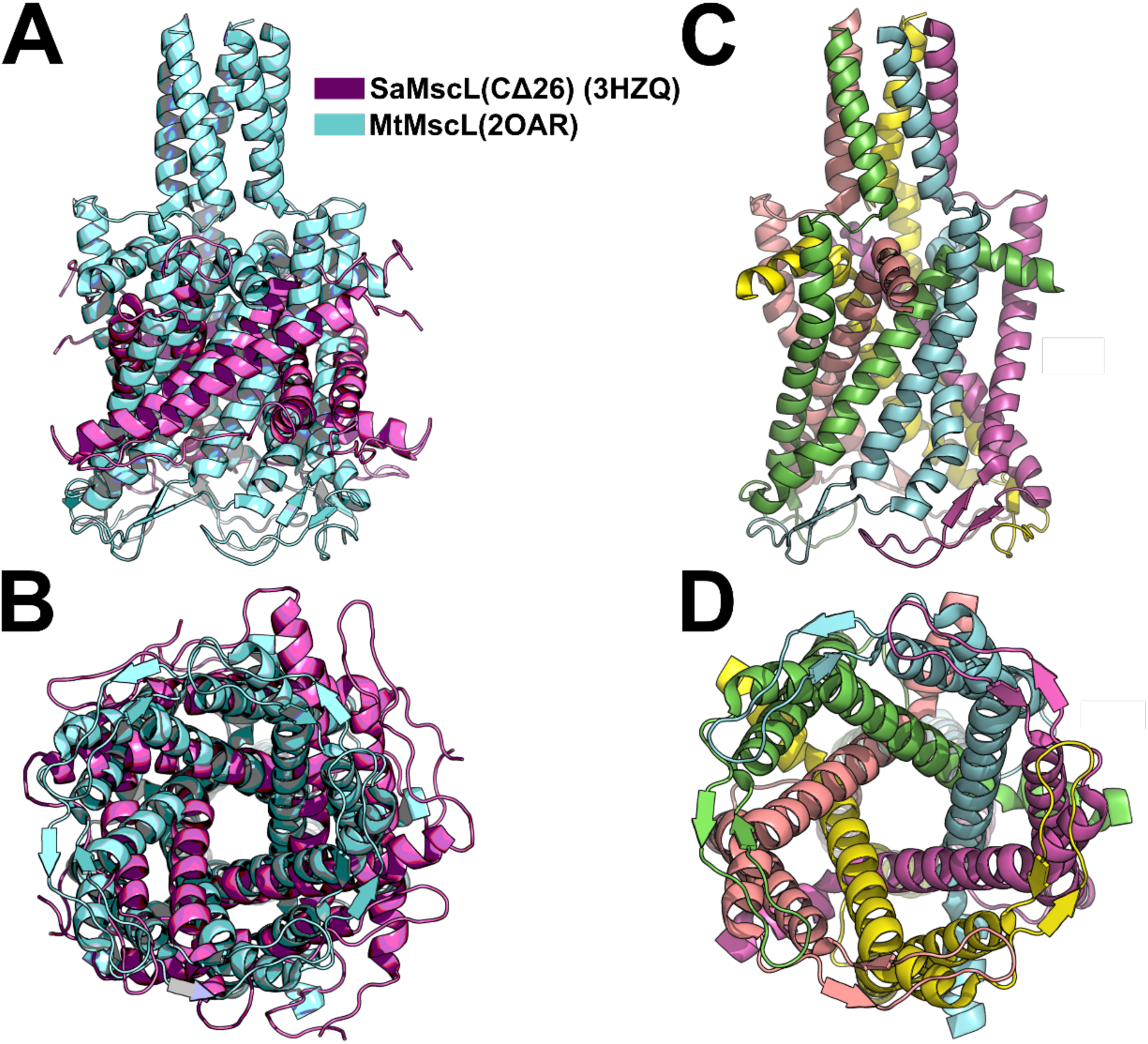
Structural alignment between the open conformation tetramer of *Staphylococcus aureus* (SaMcsLCΔ26) (3HZQ) and the pentamer of *Mycobacterium tuberculosis* (MtMscL) (2OAR) from a side and top view in A and B, respectively. C and D show the structure of the MtMscL channel, with a different color assigned to each monomer, depicted from both a side and top view, respectively. Structural alignment was performed using PyMOL (Molecular Graphics System, version 3.0, Schrödinger, LLC).

**Figure 3.**
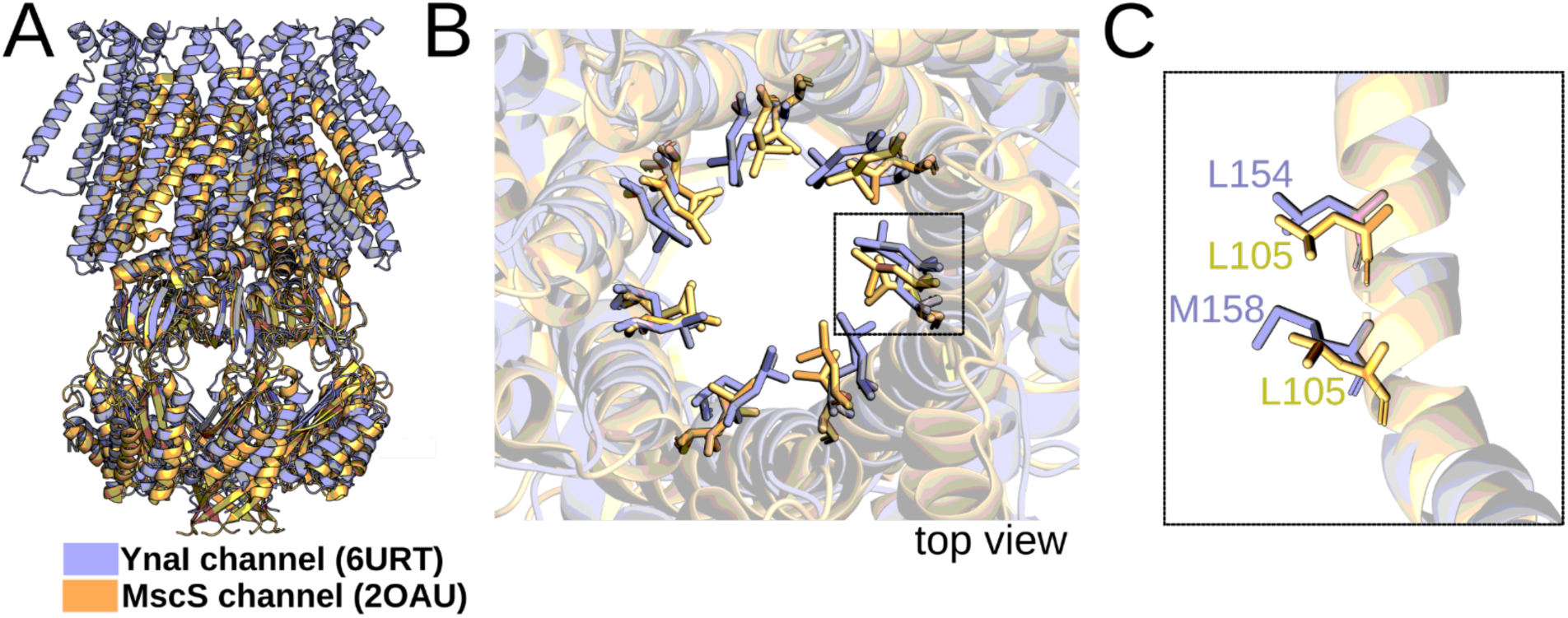
Structural alignment between YnaI and MscS channel structures of *Escherichia coli.* (A) Crystallographic structures. (B) A bottleneck formed in the pore by the analogous residues shown. Top view. (C) Side view showing the two hydrophobic layers of the bottleneck, with a focus on a single monomer. Structural alignment was performed using PyMOL (Molecular Graphics System, version 3.0, Schrödinger, LLC).

**Figure 4.**
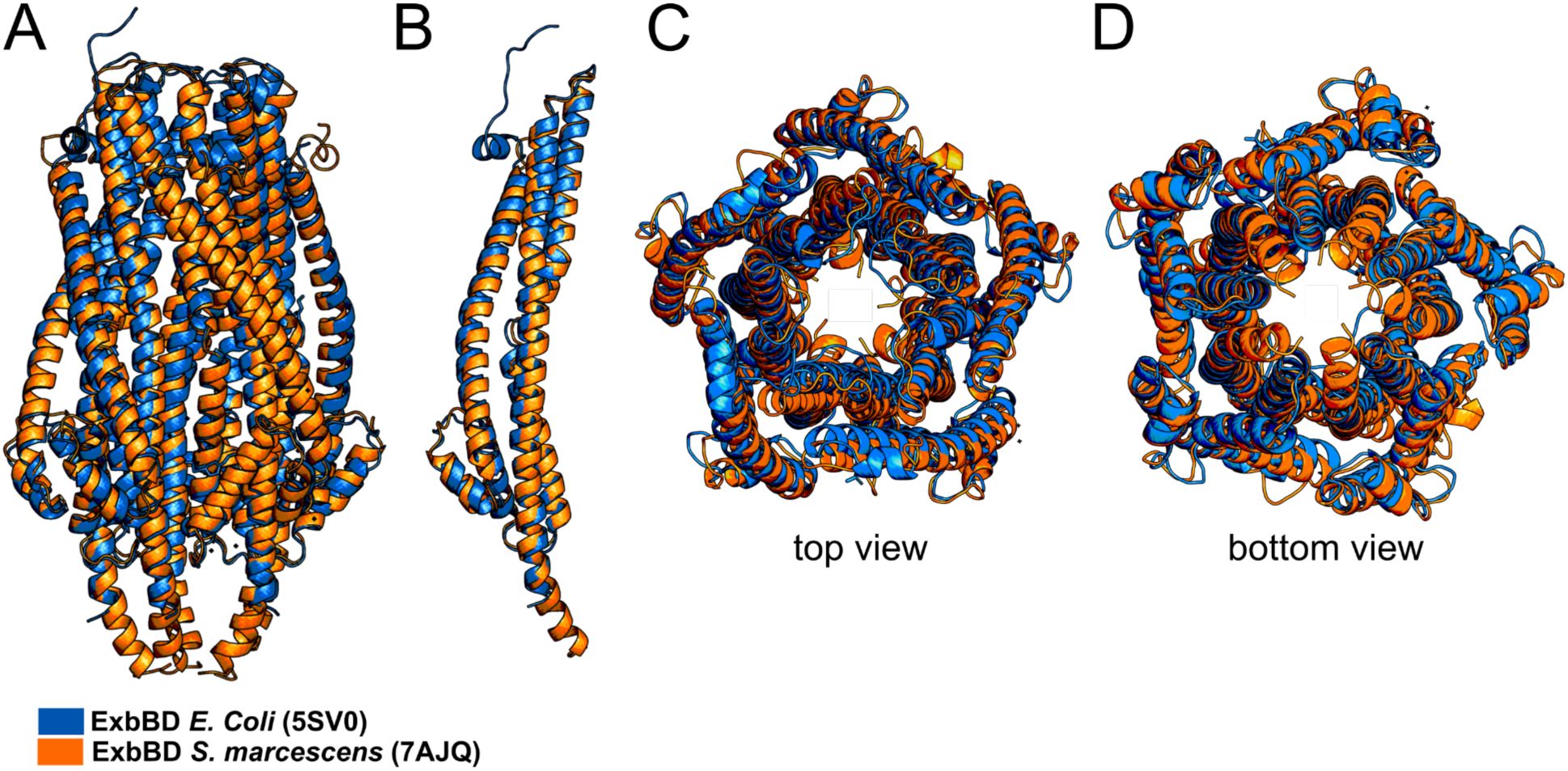
Structural alignment of the ExbB channel of *Serratia marcescens* and *Escherichia coli.* (A) Crystallographic structures. (B) One monomer from each channel. Top and bottom views of crystallographic structures in C and D, respectively. Structural alignment was performed using PyMOL (Molecular Graphics System, version 3.0, Schrödinger, LLC).

